# N440K variant of SARS-CoV-2 has Higher Infectious Fitness

**DOI:** 10.1101/2021.04.30.441434

**Authors:** Dixit Tandel, Divya Gupta, Vishal Sah, Krishnan Harinivas Harshan

## Abstract

Several variants of SARS-CoV-2 have been emerging across the globe, continuing to threaten the efforts to end COVID-19 pandemic. Recent data indicate the prevalence of variants with N440K Spike substitution in several parts of India, which is under the second wave of the pandemic. Here, we first analyze the prevalence of N440K variants within the sequences submitted from India and identify a rising trend of its spread across various clusters. We then compare the replicative fitness and infectivity of a prototype of this variant with two other previously prevalent strains. The N440K variant produced ten times higher infectious viral titers than a prevalent A2a strain, and over 1000 folds higher titers than a much less prevalent A3i strain prototype in Caco2 cells. Similar results were detected in Calu-3 cells as well, confirming the increased potency of the N440K variant. Interestingly, A3i strain showed the highest viral RNA levels, but the lowest infectious titers in the culture supernatants, indicating the absence of correlation between the RNA content and the infectivity of the sample. N440K mutation has been reported in several viral sequences across India and based on our results, we predict that the higher infectious titers achieved by N440K variant could possibly lead to its higher rate of transmission. Availability of more sequencing data in the immediate future would help understand the potential spread of this variant in more detail.

## INTRODUCTION

Past one year has witnessed the emergence of several variant strains of the SARS-CoV-2 (1). Some of them have gained advantage over the original or previously prevalent strains in spreading across populations and replacing them in due course. Of the several such variants of interest (VoI), a few have been implicated in increased infectivity and severity. By June 2020, a variant with D614G substitution had become the predominant strain over the ancestral strains (2). This variant was demonstrated to have higher fitness (3), (4) over the previously dominant ancestral strains. Later, the UK variant B.1.1.7 lineage, Brazilian variant P.1 lineage (5) and the South African B.1.351 (6) have emerged as variants of concern (VoC) (1). The newly evolving variants are expected to have better fitness over their ancestral strains. One prototype of the UK variants named 20I/501Y.V1 with B.1.1.7 lineage was demonstrated to have higher replicative fitness over an ancestral D614G strain (7). Comparative studies have also identified efficient infection and distinct pattern of cytokine induction by these three VoIs (8) indicating that each of these variants establish a unique relationship with the host.

In this study, we compared the replicative fitness and infectivity of prototypes of three SARS-CoV-2 strains. Of these strains, N440K variant with the mutation in Spike is being increasingly detected in India (https://data.ccmb.res.in/gear19/). A3i variant has a characteristic A97V substitution in RdRP (Nsp12) sequence, while A2a strain has a D614G substitution in Spike and a and P323L substitution in RdRP. N440K variant has a P323L substitution in RdRP in addition to N440K in Spike. In addition to these, each of the variant had distinct variations in other sequences. Our studies unambiguously demonstrate that N440K variant prototype has capacity to generate significantly higher titers of infectious virus in shorter duration and suggest that this feature could promote its faster spread among certain populations.

## RESULTS AND DISCUSSION

### N440K variant prevalence has been rising in certain clusters in India

We analyzed the GISAID database (9) for the prevalence of variants containing N440K substitution in Spike. A total of 1555 entries with N440K substitution could be identified from across the world. Interestingly, India contributed the largest proportion of N440K variants at 33%, followed by the USA and Germany (Figure 1A). Submissions of sequences from India contributed to 0.86% of the total submissions (Figure 1B). Of the submissions from India, 4.9% sequences contained N440K substitution (Figure 1B). However, when Indian submissions from January 2021 till 24^th^ April 2021 were analyzed, the proportion of N440K substitution significantly went up to 8.82% (Figure 1C). A further breakdown of the recent data indicates a gradual increase in the representation of this variant with March and April 2021 adding more of this variant that the previous months (Figure 1D). An increase in the proportion of N440K variant in the Indian samples is also evident with almost 10% of the sequences submitted in April 2021 carrying this substitution (Figure 1E). Importantly, higher numbers of submissions with N440K from other parts of the world were seen in the past two months, indicating its rising spread in such countries (Figure 1F). Significantly, these months also saw increased submission of sequences from India. Karnataka, Maharashtra, Telangana and Chhattisgarh together contributed to about 50% of these samples indicating the geographically localized spread of this variant in India (Figure 2 A and B). Interestingly, over 99% of the N440K variants were in the background of D641G substitution, indicating that these variants are in the lineage of A2a strain. Taken together, the data suggests that the proportion of N440K variant has been increasing gradually and suggesting its improved replicative fitness or increased infectivity.

**Figure 1.**
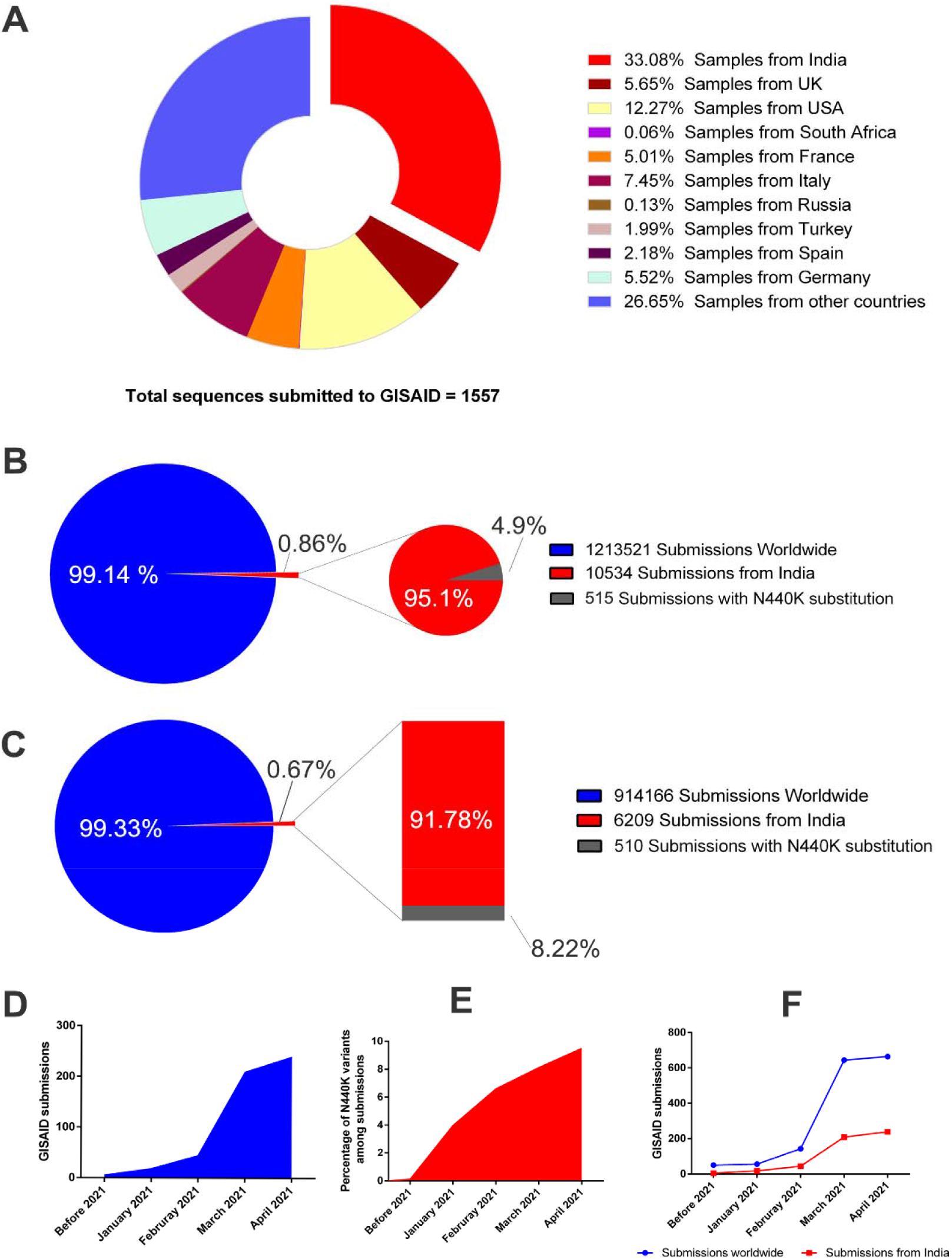
Analysis of the prevalence of N440K Spike variant of SARS-CoV-2. (A) demonstrates its prevalence across the world based on the sequences submitted to GISAID. (B) compares the proportion of N440K variant sequences submitted from India. The blue circle shows the submissions from across the world and the red circle shows the sequences from India. (C) Comparison as in (B), but the data points are selected from January, 2021 till 24^th^ April, 2021. (D) shows the increasing numbers of N440K carrying sequences from India since the beginning of 2021 and (E) its proportion in the total sequences submitted. (F) demonstrates the number of sequences containing N440K substitution from India and the rest of the world.

**Figure 2.**
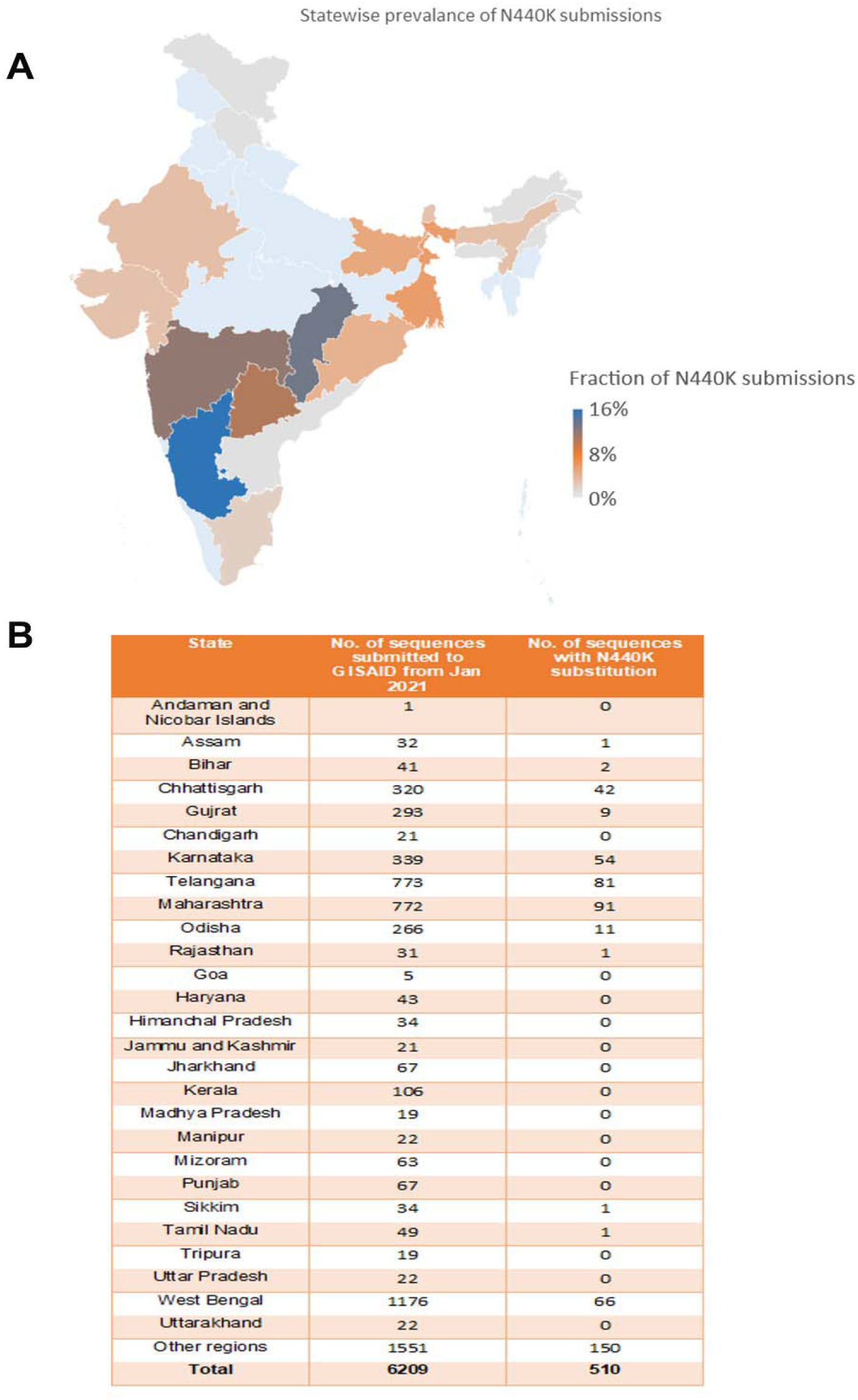
Geographic distribution of N440K variant across India as of April 24, 2021. Whole genome sequences available from GISAID were analyzed based on their sample origin. (A) demonstrates the heatmap of the distribution. The highest intensity is represented by blue while the lowest by grey. (B) Table detailing the number of sequences originated from the states and the number o N440K variants reported from the corresponding states.

### N440K mutant variant makes higher viral titers

Since the proportion of N440K variant has been rising among the sequenced samples from India, we asked if they have a growth advantage over some of the other strains. We compared the viral RNA titers of prototypes isolates of three strains from our repository. The genetic comparison is provided in Figure 3A. The growth of these viruses in cultured cells was analyzed by measuring the viral RNA titer and infectious viral titers of the supernatants. RNA titers in the viral culture supernatants are the default measure of the presence of SARS-CoV-2. Caco2 cells were infected by each of the three variants at 0.1 MOI Viral RNA in the supernatants were measured by RT-qPCR detection of SARS-CoV-2 genes at 24-, 48-, and 72-hours post-infection (hpi). As indicated by the relative RNA levels (Figure 3 B and C), the A2a strain replicated at significantly slower rate as compared to the other two strains. Both RdRP and E gene levels from A2a strain were lower than the other two by over a log. A3i strain had moderately higher levels of viral RNA than N440K strain until 48 hpi, but both attained similar RNA levels at 72 hpi. These results indicated that of the three strains, A3i has higher replicative rate than the other two while A2a replicated at the slowest rate. We repeated these experiments in another permissive cell line, Calu-3. Here again, A2a had the lowest RNA titers at 24 hpi, followed by N440K and A3i had the highest (Figure 3 E and F). However, at 48 and 72 hpi, A2a and A3i had almost similar levels of RNA while N440K had the lowest. These results indicate that A3i has the most competent replication across multiple cell lines.

**Figure 3.**
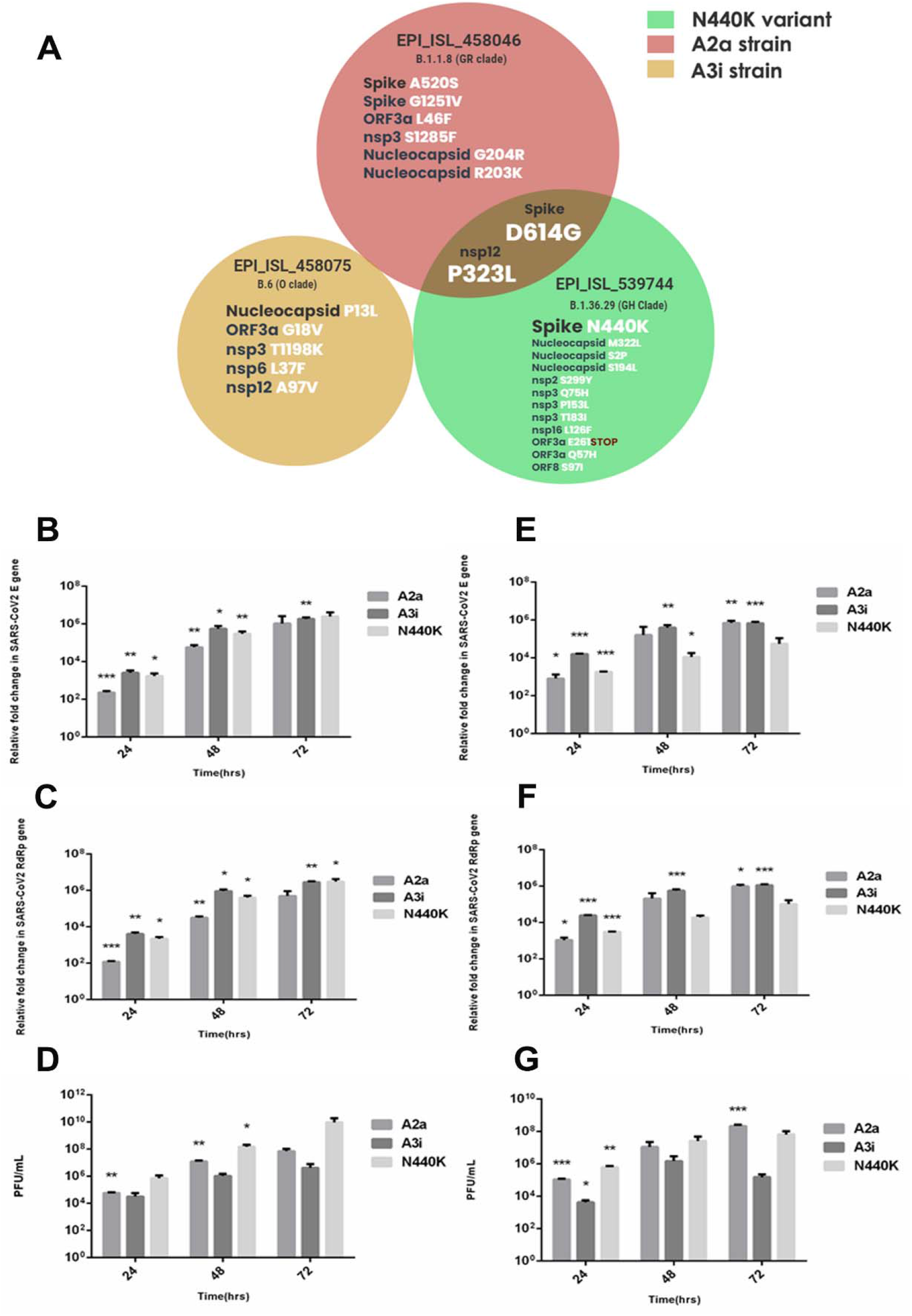
Analysis of the replicative fitness and infectious viral particle generation for three variants used in this study. (A) Comparison of genetic diversity among the three variants. (B-D) Caco2 cells were infected with 0.1 MOI of three viral variants for 24-, 48- and 72 hours. At the end of these time intervals, supernatants were collected and processed. (D) represents the absolute plaque units per mL of the supernatant collected at the respective time point. (E-G) Similar experiments and analyses performed in Calu-3 cells.

Next, we measured the infectious viral particles in the supernatants of the infected Caco2 cells. Surprisingly, N440K variant had the highest infectious viral titers from 24 to 72 hpi while A3i strain had the lowest among the three (Figure 3D). At 72 hpi, N440K variant achieved infectious titer around 10^10^ PFU/mL, which was three logs (1000 ×) higher than that of A3i. N440K variant was also produced over 10 times higher numbers than the A2a prototype. In Calu-3 cells also A3i continued to generate far fewer infectious virions (Figure 3G). N440K had higher titers at 24 hpi even though this advantage was not maintained at the later time points. These results clearly demonstrate that N440K variant prototype is able to generate far more infectious virions than both A3i and A2a strains and the A3i prototype had the lowest. Our results indicate that N440K variant has a high potential to become a dominant strain considering its capacity to produce higher titers of infectious viral particles.

N440K mutation has been reported in several clusters in India and our analysis confirms this. This mutation has been suspected to be responsible for superinfections and quick spread of the infection in certain pockets. Our studies clearly demonstrate that the variant carrying this mutation produces large titers of infectious virions more rapidly than the other two strains tested. More importantly, the ability of N440K variant to generate about 10 folds more infectious virus particles than the A2a prototype, the strain that has been in circulation worldwide, assumes significance in the context of its widespread presence in India and certain other parts of the world. N440K variant is able to generate larger amounts of viruses in shorter time and hence should be capable of rapid spread across the population. The ability to generate larger amounts of infectious virus particles in shorter amount of time could provide it significant advantage over the other competing strains in establishing itself in a large population. Their increasing proportion in certain clusters validates this observation. Whole genome sequencing data for the viral strain samples from pan-India would facilitate a better understanding of its penetration within the Indian population.

It is unclear if N440K mutation provides the virus any advantage at the entry or post-entry stages. Despite sharing the D614G variation, N440K variant differs significantly from the A2a prototype by having multiple mutations in several Nsps that are shown to interfere with the host innate immune response (10). A2a strain has been in circulation across the globe for over several months and is one of the predominant strains (11). On the other hand, A3i strain that was in circulation during the early periods of the pandemic has been very limited in circulation. One of the speculated causes for its disappearance from the population is the A97V substitution in RdRP. The dominant strains that replaced A3i have a P323L substitution that was reported to have augmented the polymerase activity, thereby enhancing their replicative fitness resulting in their dominance (12). However, our studies indicate that this strain is capable of high levels of replication, but its lower infectious titers could be responsible for its disappearance from the population. Additional mutations located in Nucleocapsid (P13L) or in the non-structural proteins could be responsible for their lower infectious titer. Our studies also underline of the fallacy of depending on RT-qPCR and highlight the importance of measuring the infectious viral titers in such comparative studies.

## MATERIALS AND METHODS

### Cell culture

Vero (CCL-81) African green monkey kidney epithelial cells, Caco2 colorectal adenocarcinoma and Calu-3 lung adenocarcinoma cells were cultured in Dulbecco's Modified Eagle's Medium (DMEM; from Gibco) with Fetal Bovine Serum (FBS; from Hyclone) and 1× penicillin-streptomycin cocktail (Gibco, 15140-122) at 37°C and 5% CO_2_. Cells were continuously passaged at 70-80% confluency and mycoplasma contamination was monitored periodically.

### SARS-CoV-2 culture, propagation and infection

Three Indian isolates of SARS-CoV-2 strains were used in this study (EPI_ISL_539744 (N440K); EPI_ISL_458046 (A2a); and EPI_ISL_458075 (A3i)). The viruses were propagated in Vero (CCL-81) cells grown in 1×DMEM deficient in serum and antibiotics. Caco2 and Calu-3 cells were infected at 0.1 MOI for 2 hours in serum-free conditions after which the media was replaced with complete media and further incubated until the time of harvesting. Supernatants collected at the end of time intervals were centrifuged to remove debris and used for RNA preparation and plaque forming assay.

### RNA preparation, RT-qPCR and plaque assay

The RNA was isolated using Nucleospin Viral RNA kit (MACHEREY-NAGEL GmbH & Co. KG) and the SARS-CoV-2 RNA was quantified using a commercial kit (LabGun™ COVID-19 RT-PCR Kit; CV9032B) in Roche LightCycler 480. The Ct values were normalized against the internal control references provided in the kit. ΔΔ-Ct values were plotted in the graph demonstrating the relative fold changes in the respective samples against the uninfected control samples. The infectious viral particle numbers in the supernatant were quantified using plaque-forming unit (PFU/mL) assay in Vero monolayers. Here, the supernatants were log diluted in 1× serum-free DMEM before infecting Vero monolayer. 2 hpi the cells were overlaid with agarose: DMEM mix containing 1% LMA and the plates were incubated for 6 days at 37°C. After the incubation, the cells were fixed with 4% formaldehyde and stained with crystal violet. The clear zones were counted and PFU was calculated as PFU/mL.

### Infographic analysis on SARS-CoV2 data

The Global initiative on sharing all influenza data (GISAID) database was used for all infographic analysis used in this study. Firstly, the common mutations in all the three strains were identified by Venny 2.1.0 (13). Following this, the Venn diagram was made using Adobe photoshop. For the graph plotting of the infographs, Graphpad Prism was used. No statistical method was applied to the graphs and no tests to attain significance was done.

### Statistical analysis

All the experiments were performed in a minimum of three independent biological replicates to generate Mean ± SEM which are plotted graphically. For statistical significance, paired end, two-tailed t-test was performed and represented as *P*-values. *, ** and *** indicate *P*-values <0.05>0.01, <0.01>0.001 and <0.001 respectively.

### Institutional ethics clearance

Institutional ethics clearance (IEC-82/2020) was obtained for the patient sample processing for virus culture.

### Institutional biosafety

Institutional biosafety clearance was obtained for the experiments pertaining to SARS-CoV-2.

## Acknowledgements

We thank the NGS team at CCMB for providing the sequences the SARS-CoV-2 strains used in this study. Several volunteers at the Centre for Cellular and Molecular Biology, who were part of COVID-19 testing, helped us gain access to the potential patient samples for virus culturing. Special thanks to Amit Kumar and Mohan Singh Moodu for their help with the logistics.

## Author Contributions

D.K., D.G., and K.H.H. conceptualized the study. D.K. set up the infections and performed RT-qPCR. D.G performed plaque forming assays. V.S performed the various comparative analyses of the viral genomes. K.H.H. wrote the manuscript with the assistance from the other authors.

## REFERENCES

1. Science Brief: Emerging SARS-CoV-2 Variants | CDC.

2. Yurkovetskiy L, Wang X, Pascal KE, Tomkins-Tinch C, Nyalile TP, Wang Y, Baum A, Diehl WE, Dauphin A, Carbone C, Veinotte K, Egri SB, Schaffner SF, Lemieux JE, Munro JB, Rafique A, Barve A, Sabeti PC, Kyratsous CA, Dudkina N V., Shen K, Luban J. 2020. Structural and Functional Analysis of the D614G SARS-CoV-2 Spike Protein Variant. Cell 183:739–751.e8.

3. Plante JA, Liu Y, Liu J, Xia H, Johnson BA, Lokugamage KG, Zhang X, Muruato AE, Zou J, Fontes-Garfias CR, Mirchandani D, Scharton D, Bilello JP, Ku Z, An Z, Kalveram B, Freiberg AN, Menachery VD, Xie X, Plante KS, Weaver SC, Shi PY. 2021. Spike mutation D614G alters SARS-CoV-2 fitness. Nature 592:116–121.

4. Korber B, Fischer WM, Gnanakaran S, Yoon H, Theiler J, Abfalterer W, Hengartner N, Giorgi EE, Bhattacharya T, Foley B, Hastie KM, Parker MD, Partridge DG, Evans CM, Freeman TM, de Silva TI, Angyal A, Brown RL, Carrilero L, Green LR, Groves DC, Johnson KJ, Keeley AJ, Lindsey BB, Parsons PJ, Raza M, Rowland-Jones S, Smith N, Tucker RM, Wang D, Wyles MD, McDanal C, Perez LG, Tang H, Moon-Walker A, Whelan SP, LaBranche CC, Saphire EO, Montefiori DC. 2020. Tracking Changes in SARS-CoV-2 Spike: Evidence that D614G Increases Infectivity of the COVID-19 Virus. Cell 182:812–827.e19.

5. Faria NR, Mellan TA, Whittaker C, Claro IM, Candido D da S, Mishra S, Crispim MAE, Sales FCS, Hawryluk I, McCrone JT, Hulswit RJG, Franco LAM, Ramundo MS, de Jesus JG, Andrade PS, Coletti TM, Ferreira GM, Silva CAM, Manuli ER, Pereira RHM, Peixoto PS, Kraemer MUG, Gaburo N, Camilo C da C, Hoeltgebaum H, Souza WM, Rocha EC, de Souza LM, de Pinho MC, Araujo LJT, Malta FS V., de Lima AB, Silva J do P, Zauli DAG, Ferreira AC de S, Schnekenberg RP, Laydon DJ, Walker PGT, Schlüter HM, dos Santos ALP, Vidal MS, Del Caro VS, Filho RMF, dos Santos HM, Aguiar RS, Proença-Modena JL, Nelson B, Hay JA, Monod M, Miscouridou X, Coupland H, Sonabend R, Vollmer M, Gandy A, Prete CA, Nascimento VH, Suchard MA, Bowden TA, Pond SLK, Wu C-H, Ratmann O, Ferguson NM, Dye C, Loman NJ, Lemey P, Rambaut A, Fraiji NA, Carvalho M do PSS, Pybus OG, Flaxman S, Bhatt S, Sabino EC. 2021. Genomics and epidemiology of the P.1 SARS-CoV-2 lineage in Manaus, Brazil. Science (80-) eabh2644.

6. Tegally H, Wilkinson E, Giovanetti M, Iranzadeh A, Fonseca V, Giandhari J, Doolabh D, Pillay S, San EJ, Msomi N, Mlisana K, von Gottberg A, Walaza S, Allam M, Ismail A, Mohale T, Glass AJ, Engelbrecht S, van Zyl G, Preiser W, Petruccione F, Sigal A, Hardie D, Marais G, Hsiao M, Korsman S, Davies MA, Tyers L, Mudau I, York D, Maslo C, Goedhals D, Abrahams S, Laguda-Akingba O, Alisoltani-Dehkordi A, Godzik A, Wibmer CK, Sewell BT, Lourenço J, Alcantara LCJ, Kosakovsky Pond SL, Weaver S, Martin D, Lessells RJ, Bhiman JN, Williamson C, de Oliveira T. 2020. Emergence and rapid spread of a new severe acute respiratory syndrome-related coronavirus 2 (SARS-CoV-2) lineage with multiple spike mutations in South Africa. medRxiv. medRxiv.

7. Touret F, Luciani L, Baronti C, Cochin M, Driouich J-S, Gilles M, Thirion L, Nougairède A, De Lamballerie X. 1207. Replicative fitness SARS-CoV-2 20I/501Y.V1 variant in a human reconstituted 1 bronchial epithelium https://doi.org/10.1101/2021.03.22.436427.

8. Abdelnabi R, Boudewijns R, Foo CS, Seldeslachts L, Zhang X, Delang L, Maes P, F Kaptein SJ, Vande Velde G, Neyts J, Dallmeier K. 2021. Comparative infectivity and pathogenesis of emerging SARS-CoV-2 variants in Syrian 1 hamsters 2 3 4. bioRxiv 2021.02.26.433062.

9. Shu Y, McCauley J. 2017. GISAID: Global initiative on sharing all influenza data – from vision to reality. Eurosurveillance.

10. Xia H, Cao Z, Xie X, Zhang X, Chen JYC, Wang H, Menachery VD, Rajsbaum R, Shi PY. 2020. Evasion of Type I Interferon by SARS-CoV-2. Cell Rep 33.

11. Baric RS. 2020. Emergence of a Highly Fit SARS-CoV-2 Variant. N Engl J Med 383:2684–2686.

12. Pachetti M, Marini B, Benedetti F, Giudici F, Mauro E, Storici P, Masciovecchio C, Angeletti S, Ciccozzi M, Gallo RC, Zella D, Ippodrino R. 2020. Emerging SARS-CoV-2 mutation hot spots include a novel RNA-dependent-RNA polymerase variant. J Transl Med 18:179.

13. -Venny-. Venn Diagrams for comparing lists. By Juan Carlos Oliveros.

